# Ambrosia beetle invasions are structured by inbreeding, intraspecific hybridisation, and bridgeheads

**DOI:** 10.64898/2026.03.31.715706

**Authors:** Thomas L Schmidt, Anandi Bierman, Elizabeth J Huisamen, John S Terblanche, Ary A Hoffmann

**Affiliations:** School of Life and Environmental Sciences, University of Sydney, NSW, Australia, 2050; Centre for Invasion Biology, Department of Conservation Ecology and Entomology, Stellenbosch University, Stellenbosch 7602, Western Cape, South Africa; Faculty of Natural Sciences, Department of Biodiversity and Conservation Biology, University of the Western Cape, Bellville, Cape Town, 7535, Western Cape, South Africa; Bio21 Institute, School of BioSciences, University of Melbourne, VIC, Australia, 3052

**Author notes:** Correspondence: Thomas L Schmidt, **Email:**.

## Abstract

When invasive populations establish in regions far from their origin, they may accumulate deleterious mutations that limit population viability and later expansion. Invasions stemming from such bridgehead populations may experience further sequential bottlenecks. However, deleterious mutations can be masked or eliminated when populations outbreed with other lineages. Here, we analyse global invasions of a species complex of persistently inbreeding ambrosia beetles, using genomic data (N=247) from invasive populations in Africa, North America and Australia, and from native populations in Asia. We mostly focus on one species of this complex (*Euwallacea fornicatus*) which poses a severe threat to tree species worldwide and is rapidly expanding its global range. We uncover a single lineage of this species across California, South Africa, and Western Australia, involving an invasive bridgehead and containing almost no nuclear genetic variation. In South Africa we identify a second lineage that has repeatedly hybridised with the first lineage. Genetic patterns in the native range indicate that such opportunistic outbreeding may be common. Despite lacking nuclear variation, the first lineage contained two CO1 haplotypes that were also observed in every hybrid lineage, pointing to heteroplasmy and possible hybrid origins of this lineage. Native populations had fewer missense mutations than invasive populations, indicating that opportunistic outbreeding may help purge fixed deleterious mutations when local lineage diversity is high. These findings highlight the importance of outbreeding even when inbreeding is common, and they demonstrate the biosecurity threat posed by subsequent gene flow into invasive populations.

## INTRODUCTION

Range expansions are biogeographical processes that shape past and present patterns of global biodiversity. Range expansions facilitated by human transportation can confer costs to the local environment or to the health of humans, livestock, or crops. These biological invasions often result in new populations far from their origin and with limited further gene flow from conspecifics. Without further gene flow, an invasive population may lack the diversity to adapt to new conditions (Smith et al. 2020) and may accumulate genetic load (Bertorelle et al. 2022), where deleterious mutations fix stochastically as purifying selection is overwhelmed by the strength of genetic drift. Genetic load can reduce establishment success (Lande 1994; Nowell Nicolle et al. 2026) and it accumulates faster when local lineage diversity is low such as when an invasive population expands into new territory (Peischl et al. 2013). The effects of this expansion load on populations are stronger and more stochastic when recombination does not occur due to inbreeding or asexuality (Peischl et al. 2015; Phillips 2025). Invasions sourced from other invasive populations (“invasive bridgeheads” (Lombaert et al. 2010)) experience these conditions to different degrees, depending on whether load has accumulated (Lombaert et al. 2025) or diminished (Facon et al. 2011) in the bridgehead population.

When deleterious alleles segregate in a population, they can be eliminated through purifying selection. Invasive populations can be effective crucibles for purging deleterious alleles, as the high levels of inbreeding in early stage or spreading invasions creates homozygotes which exposes deleterious alleles to stronger selection (Facon et al. 2011). However, once deleterious alleles fix, they can only be resegregated by reverse mutation or gene flow from lineages without the deleterious mutation, with gene flow by far the fastest option. While the above describes the situation for outbreeders, the effects of genetic load on species that routinely inbreed are less clear cut. Does inbreeding purge load so efficiently that deleterious mutations rarely fix? Or do inbreeding lineages tend to die out once sufficient load accumulates, as an analogue to Muller’s ratchet (Charlesworth and Charlesworth 1997)? And if these populations occasionally outbreed with new migrants, will this assist in purging of load?

The shot-hole borer (SHB) weevils of the *Euwallacea fornicatus* species complex present an intriguing model to explore the above issues. SHBs are haplodiploid ambrosia beetles that excavate brood galleries in trees where they establish colonies of symbiotic fungal species (predominantly *Fusarium*) for sustenance. Foundresses produce many diploid females and a few flightless, haploid males, and this brood then inbreeds repeatedly over the years that a gallery remains a viable habitat (Mendel et al. 2012; Cooperband et al. 2016; Gomez and Johnson 2019). The possibility of local outbreeding exists when males occasionally enter other brood galleries in the same tree (Gomez and Johnson 2019). SHBs are major plant pests spreading rapidly around the world by human transportation of wooden packing materials. Adults can survive in cut wood for months and continue to lay viable eggs (Eatough Jones and Paine 2015), and new invasions may in theory be initiated by a single female (Gomez and Johnson 2019).

The *E. fornicatus* species complex includes the Kuroshio shot-hole borer (KSHB), the tea shot-hole borer (TSHB), and the polyphagous shot-hole borer (PSHB) (Stouthamer et al. 2017). PSHB is considered to be the greatest biosecurity threat of this group (Hulcr et al. 2017) and can attack ≥680 host tree species and reproduce in at least 168 species (Mendel et al. 2021; Ruzzier et al. 2023). Since 2000, PSHB has spread by human transportation from its native range in Asia to every inhabited continent, with new invasions typically observed in urban centres far from other invaded regions. PSHB has established or been recently detected in locations including Southern California (c.2003 (Eskalen et al. 2012)), Hawaii (c.2006 (Rugman-Jones et al. 2020)), Israel (c.2009 (Mendel et al. 2012)), South Africa (c.2012 (Paap et al. 2018)), southwestern Australia (c.2021 (Moir et al. 2025)), Turkey (c.2024 (Acer et al. 2025)), eastern Argentina (Ceriani-Nakamurakare et al. 2023), numerous locations in Brazil (Covre et al. 2024), southern Uruguay (Ceriani-Nakamurakare et al. 2025), and southern Spain (Goldarazena et al. 2025). Other SHB have also established invasions, such as TSHB in Florida (Carrillo et al. 2016) and KSHB in San Diego, California (Cooperband et al. 2016). The damage to ornamental and horticultural trees associated with PSHB can be immense, with future economic losses in South Africa estimated at $18.45 billion over ten years (de Wit et al. 2022).

This study adopts a molecular evolutionary approach to investigate global invasions of SHB in North America, South Africa, and Australia, and native populations from Vietnam and China. We focus particularly on the South African PSHB invasion, where we identify rampant intraspecific hybridisation between two independent invasive lineages. One of these lineages was also observed in California and Australia, with no nuclear genetic variation across the three continents despite being associated with three CO1 haplotypes. We infer this lineage involves one or more invasive bridgeheads. We investigate the spread of recombinant hybrid lineages across space and time. We then look for and identify similar hybridisation patterns in the native range and infer that opportunistic outbreeding is likely common in SHB. Finally, we find evidence that invasive populations across the *E. fornicatus* species complex have more missense mutations than native populations. The findings provide a wealth of insights into SHB biology and knowledge of specific populations that can inform ongoing and future biosecurity operations.

## RESULTS

### Global SHB genomic dataset

Genomic sequence data was produced from 188 shot-hole borers (SHB; *E. fornicatus* species complex) collected from infested trees in native and invasive range areas in 2020–2021. Native range areas included Central Vietnam (n=12), North Vietnam (n=2), Fujian, China (n=1) and Hainan, China (n=1). Invasive range areas included South Africa (n=166), Los Angeles, USA (n=2), San Diego, USA (n=2), and Miami-Dade County, USA (n=2). We generated CO1 mtDNA data for all samples from the invasive range and Chinese samples, and inferred the species of each sample by matching haplotypes to those published in Stouthamer et al. (2017). All samples were identified as PSHB, except for San Diego (Kuroshio shot-hole borer) and Miami-Dade (tea shot-hole borer). Sample details are in Table S1.

In 2023–2025 an additional 59 PSHB samples were collected from across South Africa. Genomic sequence data from these samples were solely used to infer subsequent patterns. Details of these additional samples are provided in Table S1.

### Two PSHB lineages have invaded South Africa and hybridised

Initial characterisation of genetic variation within South African PSHB (n=166) revealed high levels of genetic structuring (Fig. 1a). Clustering analysis inferred seven genetic clusters characterised by high similarity within clusters and strong differentiation between them. This nuDNA structure was not echoed in mtDNA haplotype structure (Fig. 1b). Here, two mtDNA haplotypes predominated, H33 and H38 (Stouthamer et al. 2017), and these were both distributed across every nuDNA cluster. One of the nuDNA clusters, Cluster 1, contained n=119 individuals with almost no nuclear genetic variation (Fig. 1a, dotted square) but three mtDNA haplotypes (Fig. 1b). Cluster 1 was broadly distributed across South Africa as well as California and Australia (Fig. 1c), with the latter inferred by Cluster 1’s similarity to the Australian reference assembly (Bickerstaff et al. 2024) (Fig. 2b). Conversely, Clusters 2–7 were observed only within the KwaZulu-Natal region of eastern South Africa (Fig. 2c).

**Figure 1.**
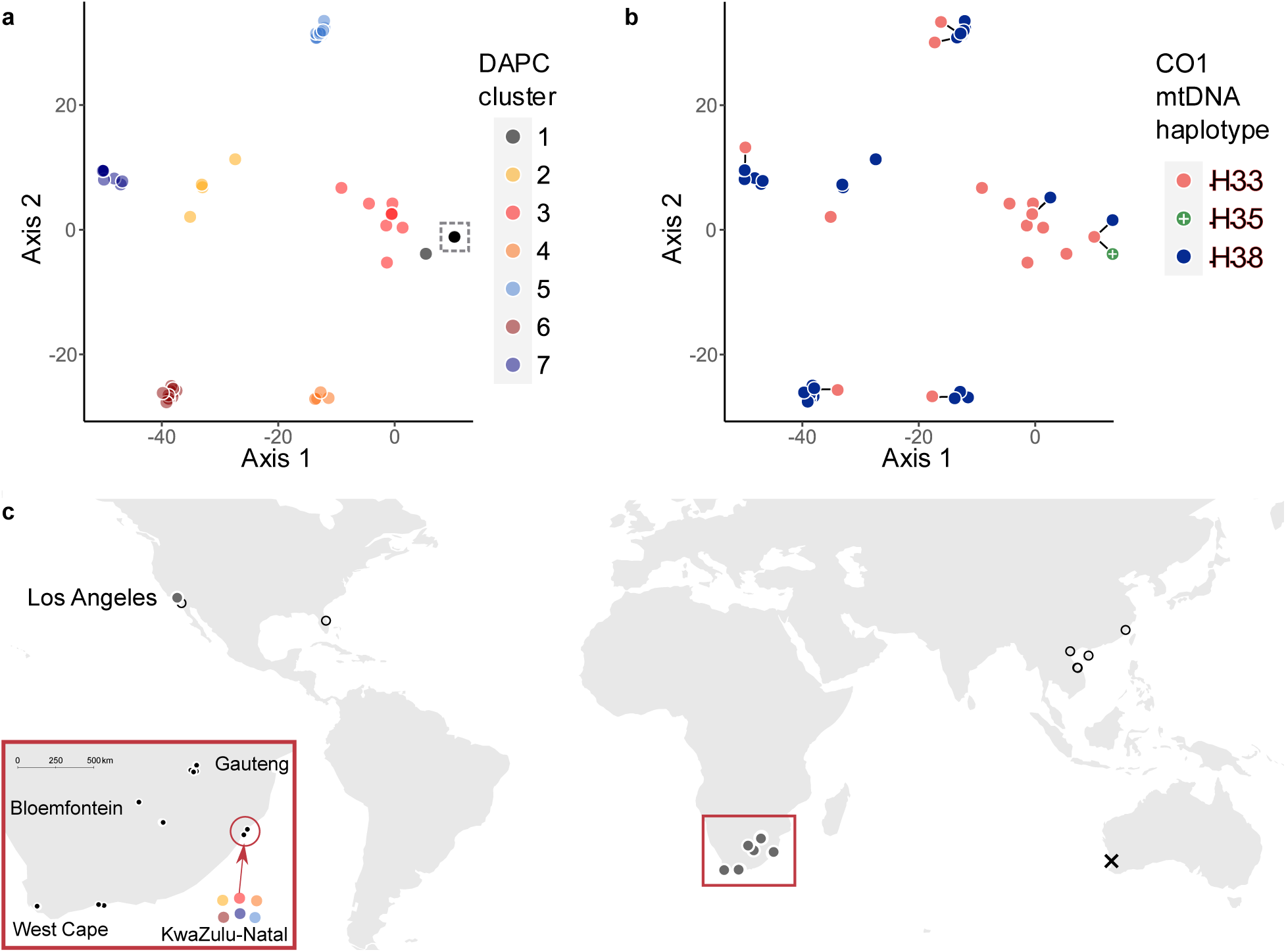
Global variation in PSHB nuDNA and mtDNA. (a) Discriminant analysis of principal components (DAPC) of South African and Californian PSHB (n=166), using K=7 clusters as inferred through *find.clusters* analysis. Six PCs were used in plotting, following guidelines (Thia 2023). Invasive lineages are Cluster 1 (n=120) and Cluster 7 (n=10). Clusters 2–6 (n=38) are hybrid clusters (see Fig. 2). The dotted square encloses n=119 Cluster 1 samples. See Fig. S1 for additional ordination analyses. (b) Identical plot to (a) but with symbology changed to reflect mtDNA CO1 haplotype. H35 haplotype was exclusive to Los Angeles. (c) Map of sample locations of all SHB. Inset details sample locations within South Africa including KwaZulu-Natal region, where Clusters 2–7 are found exclusively. Filled circles indicate DAPC cluster, including PSHB from Los Angeles (Cluster 1). Open circles indicate invasive outgroups and native range samples. Cross (×) indicates Australian reference genome location.

**Figure 2.**
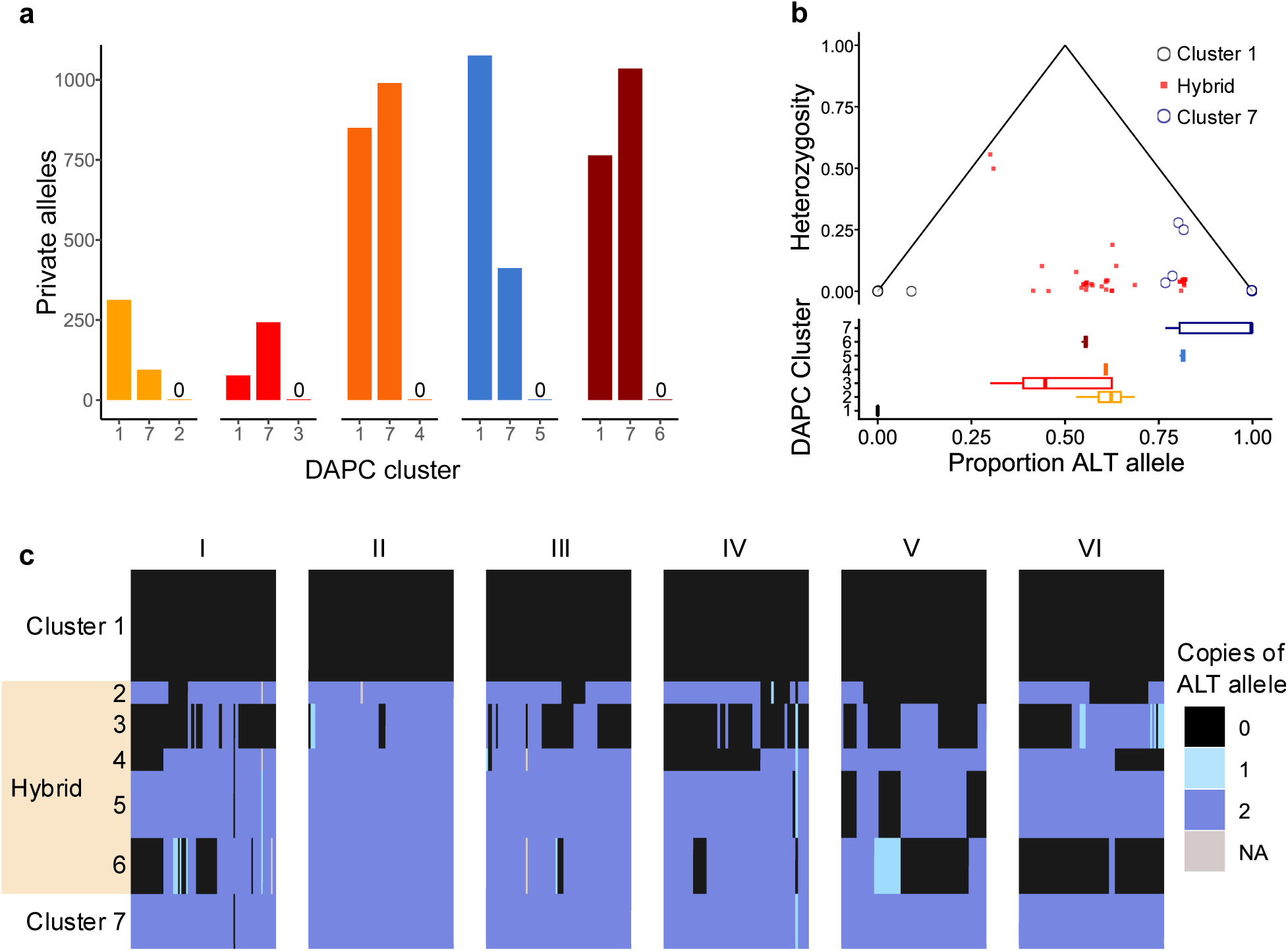
Five hybrid clusters in South Africa. (a) Private allele counts of DAPC clusters when analysed as trios. Clusters follow Fig. 1a. Each trio contains Cluster 1, Cluster 7, and one other cluster. (b) Ancestry proportion and heterozygosity of Clusters 1–7. Hybridisation triangle at the top plots individual heterozygosity at SNPs (y-axis) and proportions of alleles in each individual that are non-reference (x-axis). Horizontal boxplots at the bottom summarise allele proportions across clusters. (c) Heatmap showing PSHB genotypes across the first six chromosomes (I–VI). Individuals are sorted by cluster on the y-axis. SNP genotypes are sorted by position on the x-axis. For visualisation, individual genotypes have been replaced by the modal genotype of each cluster. Fig. S2 is the same plot as (c) with original genotypes.

We inferred that five of the nuDNA clusters (Clusters 2–6, Fig. 1a) were produced from intraspecific hybridisation between the other two (Clusters 1 and 7, Fig. 1a). This was evident from the distribution of private alleles (Fig. 2a), the proportion of non-reference alleles in each cluster (Fig. 2b), and from visual inspection of genotypes (Figs 2c, S2).

Private allele analysis of trios of populations showed that, in every trio that contained both non-hybrid clusters (Clusters 1 and 7), the remaining hybrid cluster contributed no private alleles to the total pool (Fig. 2a). Thus Clusters 2–6 contained no additional genetic variation to that observed in either Cluster 1 or Cluster 7. Clusters 2–6 likewise had intermediate proportions of non-reference alleles (x̄ = 0.589) relative to Cluster 1 (x̄ = 0.01) and Cluster 7 (x̄ = 0.846) (Fig. 2b). Together these results suggest that Clusters 2–6 each represent lineages derived from Clusters 1 and 7, either through crosses between these two clusters directly or through subsequent crosses involving other hybrid clusters.

Contrary to expectations for F1 hybrids, most samples with intermediate proportions of non-reference alleles did not have higher heterozygosity (Fig. 2b). This suggests that, if individuals from Clusters 2–6 are hybrids, they represent lineages that have hybridised several generations past and have since lost heterozygosity through subsequent inbreeding. Under this scenario, we would expect differences among the five hybrid clusters to result from clusters recombining independently at different breakpoints, as drift reduces chromosomal diversity until each cluster forms a recombinant inbred line with unique mosaic chromosomal haploblocks. We investigated this by plotting the genotypes at the first six chromosomes as heatmaps, showing the modal genotype of the cluster (Fig. 2c) and each individual genotype (Fig. S2). Plots showed that each hybrid cluster had the mosaic chromosomal structure expected from hybridisation followed by recombination and inbreeding.

As Cluster 1 was found throughout South Africa, but Cluster 7 and all hybrid clusters were found only within the KwaZulu-Natal region, we propose that Cluster 1 has invaded South Africa and spread across the country, until coming into contact in KwaZulu-Natal with a second invasion of Cluster 7. While the location where Cluster 1 initially established in South Africa is uncertain, Cluster 7 has an obvious origin in or around the KwaZulu-Natal region.

### Hybrid recombinant chromosomes reveal local movement patterns but no subsequent dispersal to other regions

Hybridisation in PSHB is likely brought about opportunistically by males entering other brood galleries in the same tree (Gomez and Johnson 2019) (Fig. 3a). PSHB has been recorded to preferentially attack tree branches already under attack by conspecifics (Mendel et al. 2017), which may help bypass tree defences but may also provide opportunities to outbreed with other brood galleries. While F1 females resulting from this hybridisation would be roughly identical, the recombinant inbred hybrids observed in KwaZulu-Natal could be produced by F1 females founding their own galleries, and recombination producing new mosaic chromosomes specific to each gallery (Figs 2c, 3a). As these galleries resume inbreeding, chromosomal diversity is rapidly lost, leading to fixation of a single recombinant chromosome within each gallery. This process explains both the diversity of hybrid clusters, the low variation within clusters, and the high differentiation between clusters (Fig. 1).

**Figure 3.**
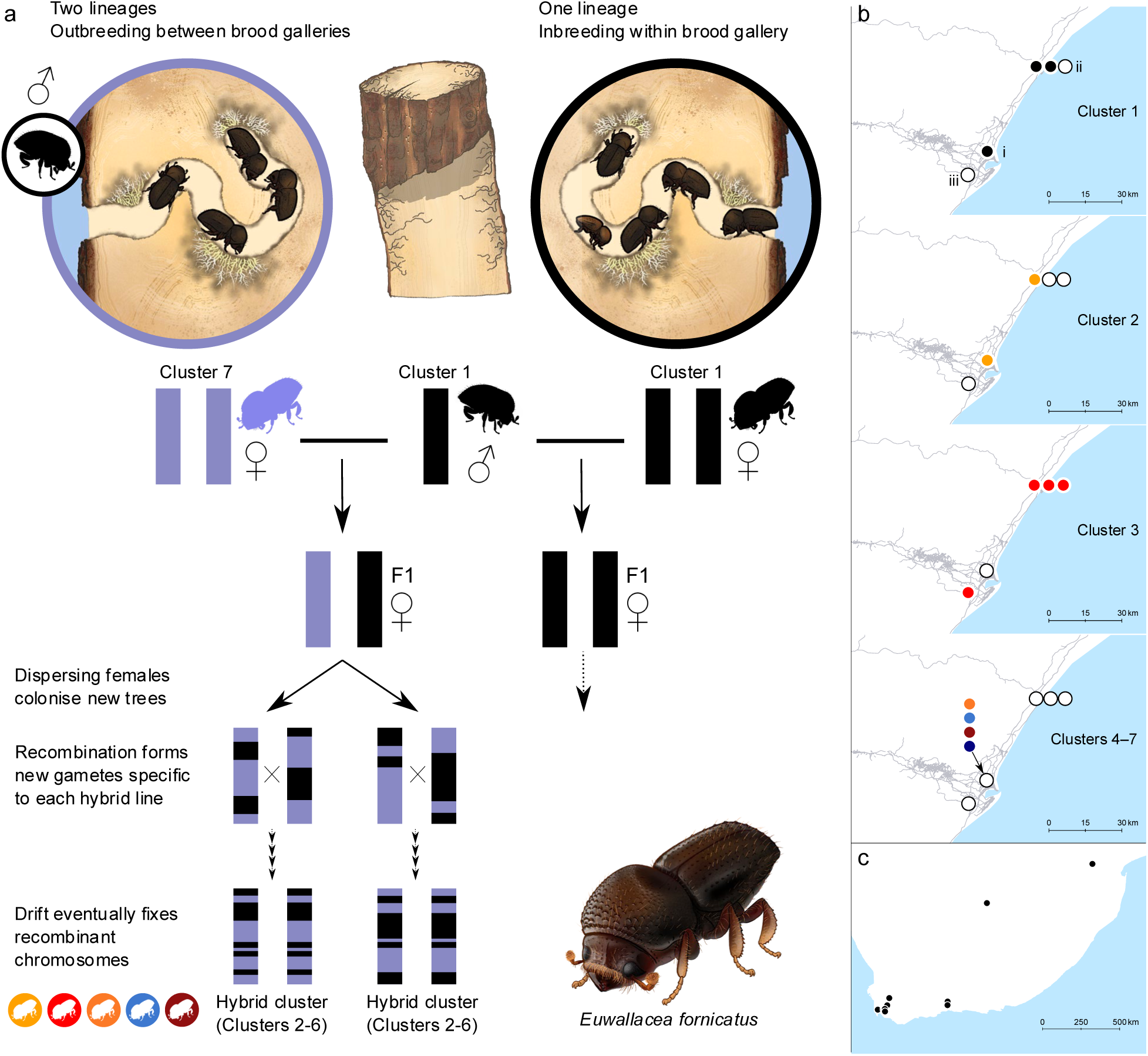
Formation and distribution of hybrid clusters. (a) Cartoon shows intergenerational outcomes of inbreeding within a single brood gallery and outbreeding among two brood galleries in the same tree. Hybrid lineages are produced from two original lineages, DAPC Clusters 1 and 7 (Fig. 1a). Members of each hybrid lineage can be identified by the unique mosaic chromosomes in each line produced by recombination and subsequent inbreeding. (b) Spatial distribution of PSHB lineages in KwaZulu-Natal province. Lineages (Clusters) follow Fig. 1a. Circles indicate locations of trees containing *E. fornicatus* (n=5 trees total), with nearby trees plotted laterally for clarity. Coloured circles indicate trees where each cluster was sampled, white circles indicate absence. Clusters 4–7 were all found in the same tree. Locations: i) Durban Botanic Gardens, ii) Simbithi Eco Estate, iii) Yellow Wood Park. (c) Locations of 59 individuals sampled at later time points, all assigned to DAPC Cluster 1 with no evidence of hybridisation (Fig. S1c).

This verbal model of SHB hybridisation requires both original lineages to co-occur within the same tree, as males are not dispersive (Gomez and Johnson 2019). Of the five trees in KwaZulu-Natal from which PSHB were sampled, one tree in Durban Botanic Gardens contained both original lineages (Clusters 1 and 7) and three of the hybrid lineages (Fig. 3b). Given the presence of these lineages, but not of all hybrid lineages, this tree likely represents one of several hybridisation sites.

While hybrid Clusters 4–6 were only detected in a single tree in Durban Botanic Gardens, hybrid Clusters 2 and 3 were detected in trees 50–60 km apart (Fig. 3b). This spatial distribution could be reached by long-distance dispersal following the fixation of recombinant chromosomes in each hybrid lineage. Having observed hybrid dispersal within KwaZulu-Natal, we then looked for evidence that Cluster 7 or its derived hybrid clusters had spread outwards into greater South Africa. For this we used the 59 samples collected in subsequent years. We found no evidence of subsequent spread, with PCA placing all 59 samples with Cluster 1 (Figs. 3c, S1c).

### South African, Californian, and Australian invasions of PSHB share a common origin involving an invasive bridgehead

We inferred evolutionary relationships between South African and non-South African SHB using a D_XY_ tree (Fig. 4). Cluster 1 formed an obvious clade with native range samples from North and Central Vietnam. Other samples from Central Vietnam clustered apart in a PCA (Fig. S1b), and formed a separate clade with Hainan, China. We refer to these two subgroups as Central Vietnam A and B respectively (Fig. 4). These occurred within ∼30 km of each other (Fig. 5c). It is possible that the Central Vietnam B lineage has established in Vietnam through long-distance dispersal from Southern China, or vice-versa. The origin of Cluster 7 was unclear, as it split from a position equidistant from Vietnam and Fujian clades. *Wolbachia* infection (strain *w*Ei) was distributed across two clades. The Hainan sample was infected (5,763 aligned reads vs x̄ = 1.1 aligned reads), as were four Central Vietnam A samples (9,595–11,719 aligned reads vs x̄ = 1.1 aligned reads). All other samples were uninfected with *Wolbachia*.

**Figure 4.**
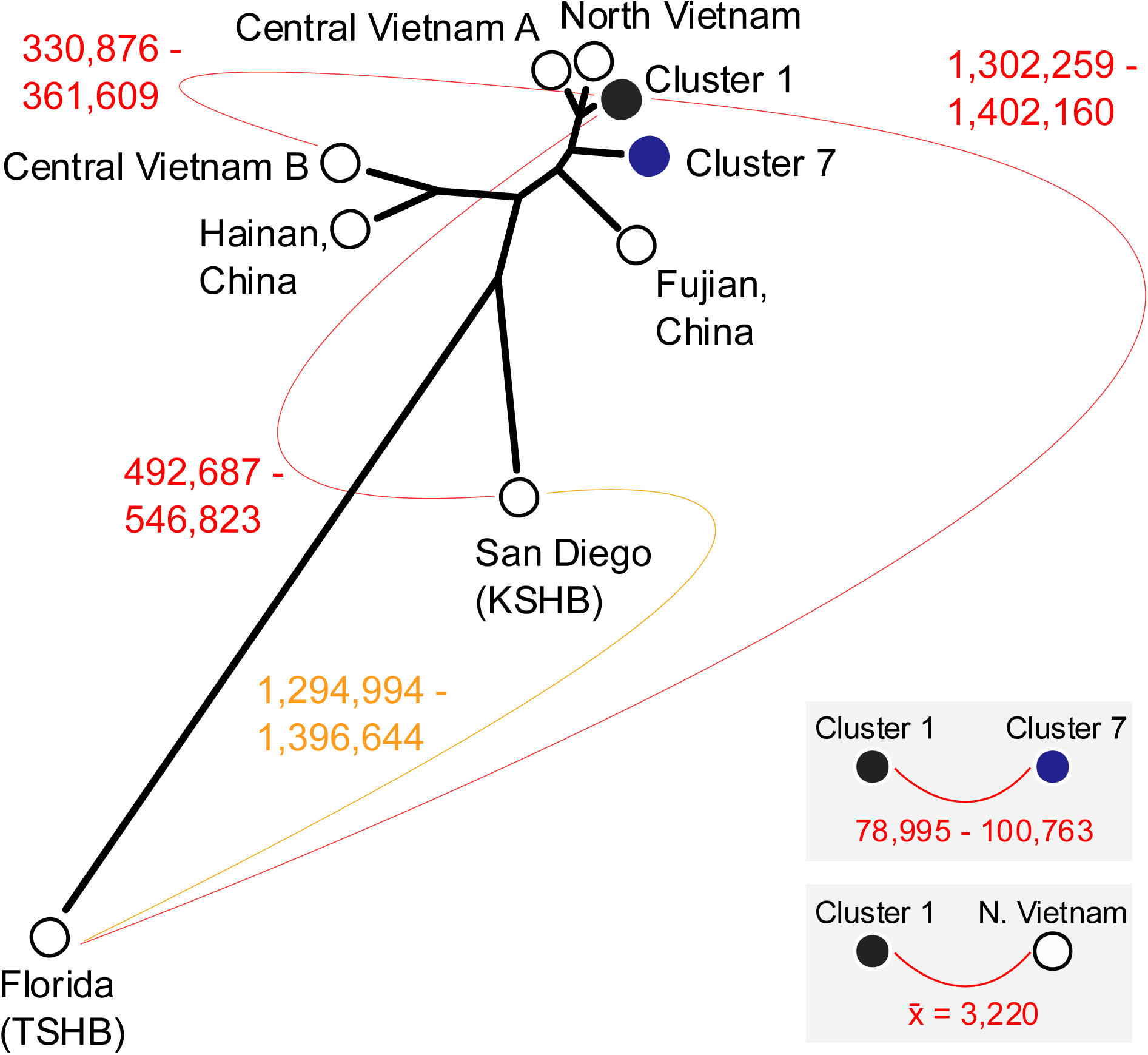
Evolutionary relationships within the SHB species complex. Tree is constructed from D_XY_ among all population pairs, inferred in 100 kb windows. Curved lines show split time estimates in units of generations, with ranges calculated from 100 bootstrap subsamples from D_XY_ windows, and using a single-nucleotide mutation rate for Colorado potato beetle of 5.8 × 10^-9^ (Xu et al. 2026).

**Figure 5.**
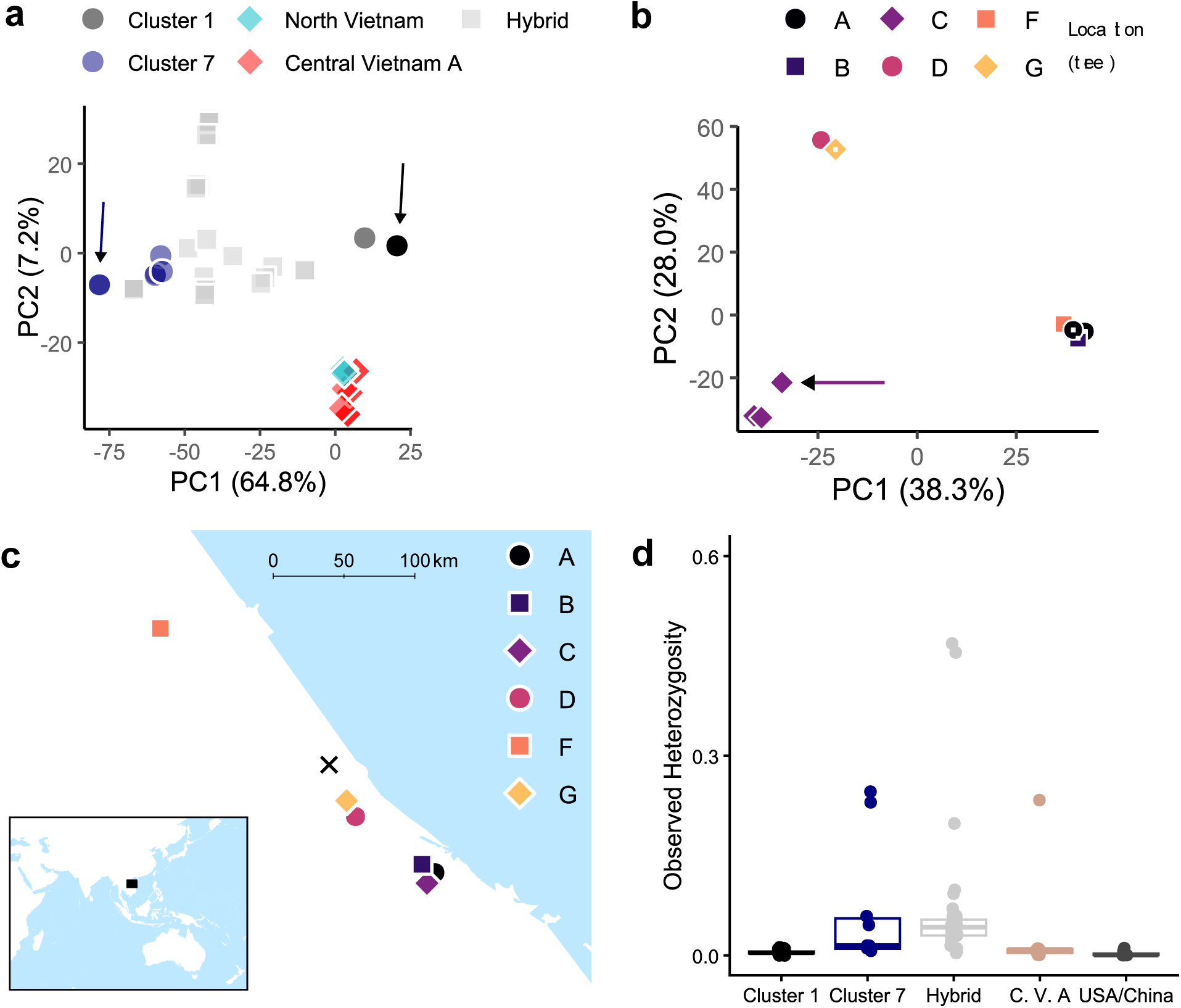
Lineage diversity and outbreeding in native and invasive range populations. (a) PCA of South Africa, Central Vietnam A and North Vietnam. Arrows point to subsets of near-identical samples in Cluster 1 (n=117, c.f. Fig. 1a) and Cluster 7 (n=6). (b) PCA of Central Vietnam A (n=9). Symbols indicate infested trees of origin. Arrow points to a single putative hybrid individual with higher heterozygosity (see d). Superimposed white squares indicate four individuals with *Wolbachia* infections. (c) Locations of the six infested trees containing Central Vietnam A samples. Symbology follows b. The cross (×) indicates the location of the Central Vietnam B samples (Fig. 4). (d) Observed heterozygosity at variable sites, filtering at MAC ≥ 3 within each of the following subsets: Cluster 1, Cluster 7, and Hybrid; Central Vietnam A (C. V. A); Fujian, Hainan, Los Angeles, Florida (TSHB), and San Diego (KSHB) (USA/China).

The age of the *E. fornicatus* species complex was estimated at ∼1.4M generations (Fig. 4). TSHB was the most evolutionarily distant species within the complex, while closer relationships were observed among PSHB, KSHB, and the Hainan/Central Vietnam B lineage that may be a fourth reproductively isolated group. To convert generation estimates to years, we assumed that eggs take 4-6 days to hatch, larvae take 16-18 days to pupate, and pupae emerge as adults after 7-9 days (Kalshoven 1958; Gomez and Johnson 2019), though development times can vary strongly with temperature (Umeda and Paine 2019). Assuming 10 gens/yr, D_XY_ results imply the species complex had a common ancestor ∼140,000 years ago.

Cluster 1 samples from South Africa were genetically indistinguishable from those from Los Angeles, USA (Fig. S1a,b) and from the Australian sample used to produce the reference assembly (Figs. 2b, S3). Thirteen Cluster 1 samples had 0 non-reference alleles at the 3060 SNPs called within South African PSHB, and both Los Angeles samples had only 4 non-reference alleles (Fig. S3). This indicates a shared evolutionary heritage of the Californian, South African, and Australian invasions. The almost total lack of differentiation between these groups suggests that they were not separately invaded from the native range, but rather that the invasive histories of these populations involves an invasive bridgehead (Lombaert et al. 2010). Such a bridgehead could be from an unsampled region in the invasive range that was invaded before or contemporaneous to California, South Africa, and Australia, or perhaps California formed a bridgehead, as this 2003 invasion was detected 9 (South Africa) to 18 (Australia) years before the others (Eskalen et al. 2012; Bierman et al. 2022; Moir et al. 2025). A combination of the above scenarios with multiple bridgeheads could also be involved.

We further investigated the native range populations in Central Vietnam A and North Vietnam (Fig. 4) to test if there was more lineage diversity in the native range than in the invaded areas. We first used PCA to investigate lineage diversity in South Africa relative to Vietnam (Fig. 5a). This showed that Cluster 1 and Cluster 7 both plotted separate to Vietnam samples and within each cluster most individuals were almost identical genetically (Fig. 5a, arrows). The remaining Cluster 1 and Cluster 7 individuals were most likely derived from hybrids that had backcrossed repeatedly to either of the original clusters. By comparison, Central Vietnam A samples showed much greater variability. A second PCA considering only Central Vietnam A samples (n=9) showed three lineages within this group (Fig. 5b). Notably, these lineages all occurred within a ∼250 km range of each other (Fig. 5c). Additionally, the trees marked A, B, and C in Fig. 5c were all sampled from the same location but contained two distinct lineages (Fig. 5b). Such a high level of lineage diversity in the native range makes it unlikely that the PSHB invasions of California, South Africa, and Australia were independently sourced from the native range, although the California invasion might yet have been sourced directly from the native range to act as a bridgehead population for South Africa and Australia.

### Hybridisation appears common in SHB

The hybridisation observed in the invasive South African range raises the possibility that similar processes may also occur in the native range. We investigated this in the Central Vietnam A samples (Fig. 5b,c). These had two characteristics conducive to opportunistic outbreeding or hybridisation. First, as two distinct lineages were observed in the co-located trees A, B, and C (Fig. 5b,c), dispersal by flight would likely be sufficient for these lineages to invade the same tree and create outbreeding opportunities. Second, a single lineage (Fig. 5b, bottom right) was observed across two locations ∼250 km apart (Fig. 5c, trees A, B, and F). This shows long-distance dispersal, which redistributes lineages across great distances to allow future outbreeding events to involve differentiated lineages. Finally, although four Central Vietnam A samples carried *Wolbachia* infections, these were distributed across two lineages and each lineage also had uninfected individuals (Fig. 5b), strongly suggesting that these SHB populations did not experience *Wolbachia*-induced reproductive isolation such as through bidirectional cytoplasmic incompatibility (Hoffmann et al. 1990).

Given this potential for hybridisation in the native range, we looked for patterns of individual heterozygosity that could reflect recent outbreeding. We observed individuals with much higher heterozygosity at variable sites in Cluster 7, Hybrid clusters, and Central Vietnam A (Fig. 5d). This points to one of the nine Central Vietnam A individuals having a recent history of outbreeding. This individual is indicated by the red arrow in Fig. 5b, and was sampled from infested tree C which was co-located with trees A and B that shared a distinct genetic background from C (Fig. 5b).

### Mito-nuclear discordance is likely due to heteroplasmy, not reciprocal backcrossing

Although Cluster1 had almost no nuDNA variation, it contained three mtDNA haplotypes, two of which (H33 and H38) were distributed across all original and hybrid clusters in South Africa (Fig. 1b). H33 was the most common haplotype in Cluster 1 (93%, 110 of 118 South African individuals), while H38 was the most common in Cluster 7 (90%, 9 of 10 individuals). Mito-nuclear discordance is frequently observed in pests (Sun et al. 2015; Schmidt et al. 2025) and here we consider two hypotheses for why this might be occurring in PSHB. First, it could be caused by repeated cycles of inbreeding and hybridisation, as noted in studies of other bark beetles (Jordal et al. 2002). Under this hypothesis, H33 was introduced by Cluster 1, and H38 by Cluster 7, and mito-nuclear discordance has resulted from backcrossing following hybridisation. Alternatively, these patterns could be caused by heteroplasmy, as recently reported in Californian PSHB populations that may correspond to our Cluster 1 (Rugman-Jones et al. 2024).

Two observations suggest that heteroplasmy is the cause of mito-nuclear discordance here. The first is that both haplotypes were found in every hybrid cluster. Assuming that our model for how hybrid clusters form (Fig. 3a) is correct, it would be highly unlikely that two hybrid lineages, started by F1 females with different mtDNA haplotypes, would attain the same nuclear background through backcrossing with the same male, particularly in every instance of hybridisation. The second is that we observed H38 in Cluster 1 individuals far from the hybridisation region in KwaZulu-Natal (Fig. S4). If H38 was introduced by Cluster 7 and backcrossed into Cluster 1, it appears unlikely that only this backcrossed background would reach the Western Cape region without additional movement of other lineages, even in samples taken years afterwards (Fig. 3c).

### Hybridisation may help purge deleterious mutations

One explanation for why opportunistic outbreeding occurs in SHB is that it provides a means to purge deleterious mutations that have fixed in some lineages but not in others. F1 hybrids of lineages fixed alternately at the deleterious and non-deleterious alleles at these sites will be heterozygous, and as these hybrid lineages return to inbreeding they will fix one of the alleles (Figs 2c, 3a), with the likelihood of fixing the non-deleterious allele proportional to the strength of purifying selection relative to drift. Hybridisation also produces new combinations of alleles through recombination (Figs 2c, 3a) which may vary in their fitness effects, increasing the impacts of natural selection on SHB lineages.

For hybridisation to purge deleterious mutations, there must be sufficient local lineage diversity for deleterious mutations to resegregate. Invasive lineages that establish far from their origin are unlikely to have this lineage diversity and thus all outbreeding will be with lineages fixed at the same deleterious alleles. As native range populations have greater local lineage diversity (Fig. 5), we hypothesised that they will have fewer deleterious mutations than invasive range populations, as a greater proportion of these mutations will be open to purging through outbreeding.

To investigate the potential effects of hybridisation of deleterious mutations, we first compared the distribution of synonymous and nonsynonymous variants that were segregating within native range lineages of PSHB (North Vietnam and Central Vietnam A; Fig. 4) with those segregating within invasive range lineages (Cluster 1 and Cluster 7 of South Africa). For this, we processed native range and invasive range populations separately, to retain biallelic SNPs of which the vast majority were exclusively found in homozygous form (see Materials and Methods for filtering details). We assumed that, for any given SNP, the two alleles would either both have the same fitness or one allele would be deleterious. We also assumed that synonymous substitutions would always result in two alleles with the same fitness, while nonsynonymous substitutions would be on average deleterious. We used the Ensembl Variant Effect Predictor (McLaren et al. 2016) to predict the effects of each variant in the coding sequence based on its codon position. Although more sophisticated methods using the site frequency spectrum can infer the distribution of fitness effects across variants (Sendrowski and Bataillon 2024), the persistent inbreeding in SHB prevents these methods being applied in these systems due to severe distortions in the site frequency spectrum. As our analysis is within relatively short evolutionary timescales, we ignored the possibility of multiple substitutions at a single site.

In native range populations, we detected 23 nonsynonymous and 157 synonymous variants (dN/dS = 0.146) (Fig. 6a). In invasive range populations, we detected 54 nonsynonymous and 280 synonymous variants (dN/dS = 0.193). These results point to purifying selection operating in native and invasive lineages, as dN/dS is << 1. The 24.4% lower dN/dS in native lineages suggests a potential role of hybridisation for purging mutations, though the statistical power is low owing to small numbers of SNPs (two-tailed binomial test, expected p = 0.1278, 54 successes in 334 trials, p = 0.0709).

**Figure 6.**
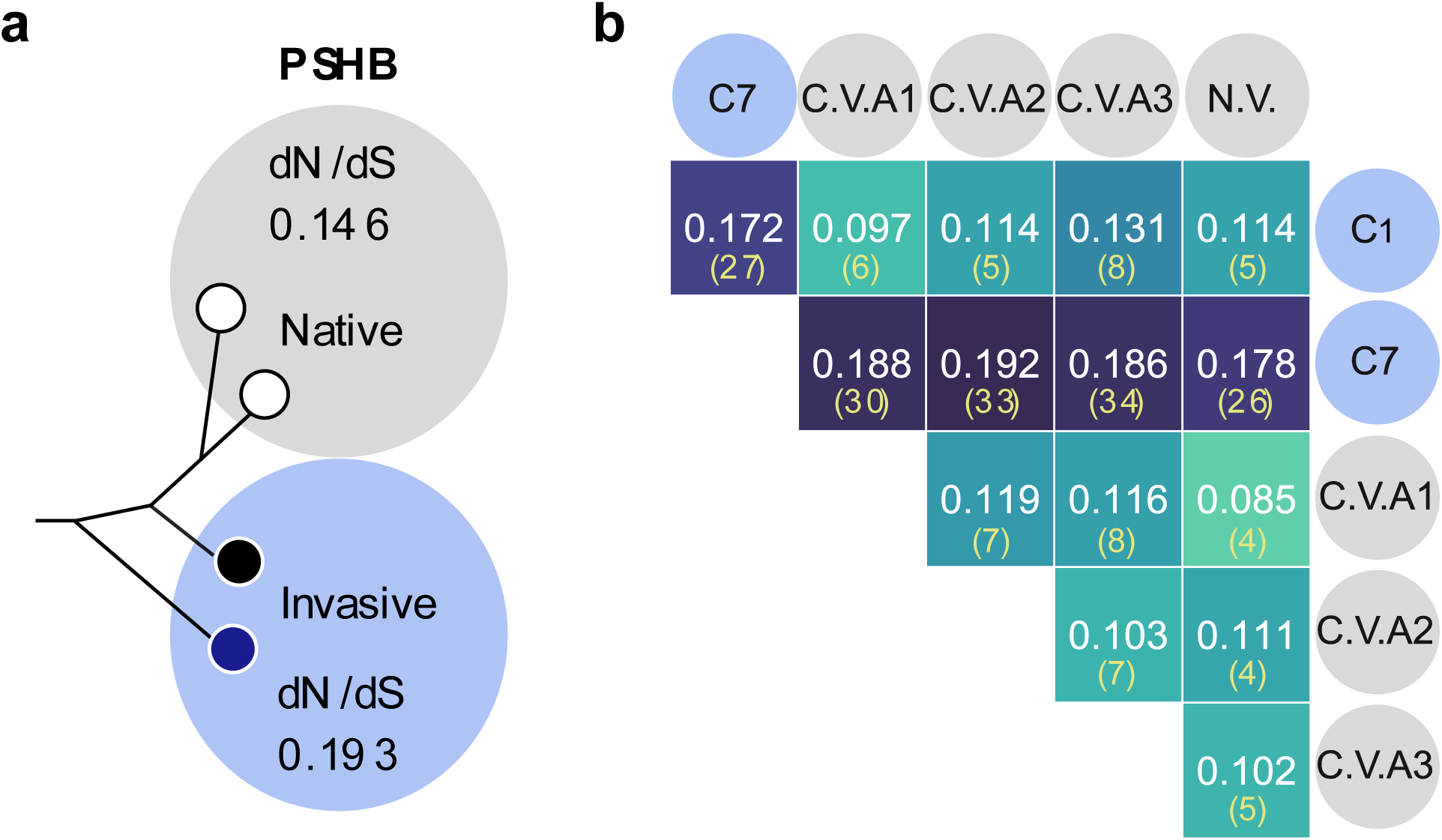
Ratios of non-synonymous and synonymous SNPs (dN/dS) within and between lineages. (a) dN/dS within native and invasive PSHB lineages. The analysis compares dN/dS at SNPs segregating among North Vietnam and Central Vietnam A (grey circle) and among Cluster 1 and Cluster 7 (blue circle). (b) dN/dS of SNPs fixed for different alleles (F_ST_ = 1) between native and invasive PSHB. White text and cell colour indicate dN/dS, yellow text in parentheses shows number of nonsynonymous substitutions. C1/C7 = Cluster 1/7; C. V. A = Central Vietnam A; N. V. = North Vietnam. C. V. A suffix numerals indicate the three lineages in Central Vietnam (Fig. 5b), analysed separately. Only samples with no evidence of admixture were included (Fig. 5a,b).

Having observed a greater number of missense mutations in invasive range populations, we investigated dN/dS for the branches specific to each population. For this, we focused on fixed differences (F_ST_ = 1) between each population pair from Vietnam and South Africa, omitting admixed individuals identified through PCA (Fig. 5a,b). This analysis showed that the majority of nonsynonymous substitutions in the invasive range were from Cluster 7, while Cluster 1 had proportions more similar to the native range populations (Fig. 6b).

## DISCUSSION

Molecular evolutionary studies of wild populations typically aim to connect patterns of DNA sequence variation to predictions from evolutionary theory. For organisms with cryptic or enigmatic life-histories, such studies can also help reveal aspects of the organism’s biology that are difficult to identify with other approaches. This study has used population genomic approaches to uncover a range of insights into the globally invasive *E. fornicatus* species complex. For PSHB, we uncovered two invasive lineages. The first was likely derived from a bridgehead, and was observed in Southern California (invaded c.2003 (Eskalen et al. 2012)), South Africa (c.2012 (Paap et al. 2018)), and southwestern Australia (c.2021 (Moir et al. 2025)). This lineage contained almost no nuclear genetic variation but three segregating mitochondrial backgrounds. The second lineage was observed only in eastern South Africa, where it has hybridised repeatedly with the first lineage to produce a series of recombinant inbred lines. We show that the lineage diversity required for hybridisation is also present in native range populations in Central Vietnam, and we found evidence of native range hybridisation, suggesting that this may be common in SHB populations. We hypothesised that hybridisation may help purge deleterious alleles that are fixed in persistently inbreeding lineages, and that invasive populations lacking local lineage diversity would not benefit from this process. Consistent with this hypothesis, we found evidence that native range populations had fewer variants at non-synonymous sites than invasive range populations.

The SHB populations considered here present an interesting case study in invasion biology, as their life histories are shaped by both inbreeding and hybridisation. Species for which inbreeding is obligate can appear as well-suited to invasion as they suffer fewer additional inbreeding consequences (i.e., genetic load) relative to that experienced in their native range (Warren and Mokadam 2025). On the other hand, without recombination through hybridisation, genetic load in an expanding invasion of inbreeders leads to highly stochastic dynamics where local populations repeatedly go extinct (Peischl et al. 2015; Phillips 2025). Our findings here show purifying selection in all SHB populations, even those most isolated from conspecifics. Native range populations had comparable dN/dS to various outbred taxa (Weber et al. 2014; Figuet et al. 2016), though lower dN/dS ratios of 0.02–0.05 have been recorded in other insects (Cunningham et al. 2015; Tierney et al. 2015).

In SHB, purifying selection operates within brood galleries, where *de novo* deleterious mutations will rapidly be eliminated or else fixed through drift. In the latter case, outbreeding with other lineages provides additional opportunities to mask and eliminate these mutations, though following outbreeding the mutations may still become fixed through drift. When local lineage diversity is low, drift may fix a deleterious mutation across all lineages, at which point it can only be eliminated by *de novo* mutation or (more likely) gene flow. Our results showing the invasive Cluster 7 lineage had higher branch-specific proportions of deleterious mutations than native range lineages support this purifying selection model. Note that strongly deleterious mutations are expected to be eliminated quickly under this model as these will be persistently exposed to selection as homozygotes (Glémin 2003). Instead, only deleterious mutations with small effects on fitness are expected to become fixed, though their effects may be cumulative (Charlesworth and Charlesworth 1997). The above results accord with general expectations that under strong purifying selection, dN/dS will be higher when effective population size (N_e_) is small (Subramanian 2013). Although N_e_ within brood galleries may be similar in native and invasive lineages, the N_e_ of the wider population will be higher in the native range due to more lineage diversity.

For invasive populations where outbreeding is the norm, purging of deleterious mutations can occur through invasion bottlenecks and will target strongly deleterious, recessive mutations (Glémin 2003; Facon et al. 2011). Whether purging is successful appears highly variable (Glémin 2003; Facon et al. 2011; Lombaert et al. 2025). Purging requires bottlenecks of moderate size (e.g., 40-300 individuals (Glémin 2003; Facon et al. 2011)), and invasions from small propagules may not be observed if they fail to establish (though see (Schmidt et al. 2023)). In contrast, SHB may be able to establish an intercontinental invasion from the long-distance dispersal of a single female, with subsequent long-distance movements resulting in SHB rapidly spreading to distant areas as observed in KwaZulu-Natal hybrids (Fig. 3b) and in recent expansions between South American cities (Ceriani-Nakamurakare et al. 2023; Covre et al. 2024; Ceriani-Nakamurakare et al. 2025). Non-lethal deleterious mutations might eventually be expected to become fixed and affect population viability as each new invasion accumulates more load (Peischl et al. 2013; Phillips 2025). However ‘genetic invasions’ (Schmidt, Endersby-Harshman, et al. 2021) of new SHB lineages into already invaded regions may reduce a fixed load through hybridisation.

Clusters 1 and 7 in KwaZulu-Natal, South Africa, represents one such genetic invasion. The hybridisation there is likely facilitated by PSHB preferentially attacking trees and branches already under attack by conspecifics (Mendel et al. 2017), though it is unclear whether this behaviour evolved to promote outbreeding or to bypass tree defences. Cycles of hybridisation and inbreeding can rapidly produce new lineages of distinct genetic character (Figs 1–3). Indeed, Cluster 1 — the lineage found in Los Angeles, South Africa, and Western Australia — may be the product of recent hybridisation. This is consistent with the observed mtDNA diversity between these locations despite a lack of nuDNA diversity, most likely due to heteroplasmy. Heteroplasmy was recently observed in Californian PSHB (Rugman-Jones et al. 2024), and was found to be stable across at least ten generations. Here, the presence of H38 in Cluster 1 individuals in the Western Cape region, far from the other lineages in the east, suggests that H33 and H38 were both present in Cluster 1 before it invaded South Africa. This in turn points to heteroplasmy being stable over even longer periods of time than observed in California. Moreover, while the second haplotype in California (H35) may be recently derived from H33, and thus could result from acquisition of new mutations (Rugman-Jones et al. 2024), in South Africa the two haplotypes are distantly related (Lantschner et al. 2026). The heteroplasmy observed in this study has therefore been produced by another mechanism, such as paternal leakage during intraspecific hybridisation (Schwartz 2021).

We associated Cluster 1 with an invasive bridgehead which may have been produced from the recombination of multiple nuclear lineages. Further evidence of the same bridgehead involving California, South Africa, and Australia comes from the recent report of a fungal associate of PSHB, *Graphium euwallaceae*, detected in the Australian PSHB invasion being genetically linked to both California and Vietnam (Moir et al. 2025). If Cluster 1 was formed through hybridisation, with outbreeding in its recent history allowing it to purge *de novo* mutations, this could potentially explain why branch-specific dN/dS analyses involving Cluster 1 indicated fewer nonsynonymous substitutions than similar analyses involving Cluster 7. However, our dN/dS analyses are based on a small number of SNPs and rely on assumptions around the selection coefficients of missense mutations. As such, specific inferences around genetic load and the purging of mutations will need further analysis involving whole genomes and phenotypic data. Similarly, greater sampling of the native range will help clarify whether the patterns of high lineage diversity we observed in Vietnam are found in other areas, and directed sampling near ports may identify if the hybrid bridgehead inferred for Cluster 1 has a specific origin linked to trade routes.

Given that outbreeding occurs in invasive (Fig. 2) and native range (Fig. 5) populations, we can also identify lineages that do not hybridise despite being in proximity. Central Vietnam A and B (Fig. 4) are two such lineages, strongly genetically differentiated and from the same location in Vietnam. Perhaps these lineages are reproductively isolated though it is also possible that Central Vietnam B may have very recently become established in Central Vietnam from an origin closer to Hainan, China (Fig. 4). It is possible that there are additional reproductively isolated lineages in the *E. fornicatus* species complex, given the broad native geographical range. The behaviour of invasive lineages that come in close contact following long-distance dispersal may represent a means of identifying reproductive isolation and is also important from an applied perspective given that members of the *E. fornicatus* species complex are thought to differ in virulence (Hulcr et al. 2017). Exploring SHB lineage associations with fungi including *Fusarium* and *Graphium* will also be vital given potential fungal effects on host virulence and polyphagy.

Results from this study raise important issues for biosecurity and pest management of SHBs. Firstly, the risk of genetic invasions is clear from the observed hybridisation and the evidence of more missense mutations in one of the invasive lineages. This demonstrates the importance of ongoing border biosecurity to intercept and reduce propagule pressure of new incursions even after a species establishes, given that such incursions may increase population fitness. While genetic invasion monitoring is frequently done to prevent the incursion of positively selected pesticide resistance alleles into new areas (Schmidt et al. 2019), such monitoring in SHB would aim to prevent masking and elimination of deleterious alleles. The introduction of novel genetic diversity is an evolutionary analogue of increasing propagule pressure (Lockwood et al. 2005), though it remains to be seen whether a lack of gene flow compromises invasive SHB populations. Secondly, the presence of three CO1 mtDNA barcodes (H33, H35 & H38) within the homogenous nuclear lineage of Cluster 1 suggests that CO1 barcodes are not particularly useful for differentiating SHB below the species level. Thirdly, we have shown how the unique genetic structure of each hybrid lineage makes spreading lineages easy to track genetically (Fig. 3b). This will remain the case for all new brood galleries established from a lineage that have not subsequently outbred. Thus, while mtDNA may lack power for tracking local lineages, nuclear amplicons developed around local variation could be useful for this purpose and also provide insights into the frequency of outbreeding.

## MATERIALS AND METHODS

### Sampling

South African PSHB were collected from 24 locations across five provinces between June 2020 and April 2021. Locations were differentiated by geographic barriers (e.g. breaks in green space or highways). Samples were collected from 23 different tree species, including those indigenous and alien to South Africa (Table S1). Approval for sampling from botanical gardens and national parks was obtained through the relevant permits for those locations, and approvals for handling and transport of dead specimens were obtained from the Department of Agriculture Forestry and Fisheries, South Africa. In addition to samples from South Africa, PSHB were obtained from the putative native range in China (Fujian, Fuzhou and Hainan, Haikou) and Vietnam (central Vietnam and north Vietnam) and from an invasive population in Los Angeles, USA (Table S1). Additional samples from the *E. fornicatus* species complex were obtained from San Diego, USA (Kuroshio shot-hole borer), and Florida, USA (tea shot-hole borer). In 2023–2025, an additional 59 samples were collected from 26 additional locations in South Africa. All sample details are listed in Table S1.

Infested branches were removed from the tree host and placed in 10 L plastic sealed containers for transport to the laboratory. Branches were then split to verify gallery formation and find adult beetles, eggs, larvae and pupae (Paap et al. 2018). Beetles were collected from exposed galleries with a needle or paintbrush, then placed in 1.5 mL microcentrifuge tubes (Eppendorf, Germany) filled with absolute ethanol and stored at −20 °C.

### DNA extraction and sequencing

Genomic sequence data were produced using a reduced representation DArT-Seq^TM^ workflow (Kilian et al. 2012). Heads and thorax were dissected from *E. fornicatus* specimens and shipped to Diversity Arrays Pty Ltd (Canberra, Australia) for DNA extraction and genotyping. Detailed methodologies for DArT-Seq^TM^ DNA extraction and sequencing are explained in (Kilian et al. 2012). Sequencing was conducted on the Illumina HiSeq2000, generating approximately 2.5 million sequences per sample. Raw sequence reads were output in fastq format and used in all downstream analyses. Some samples were used in multiple runs; these fastq files were concatenated into single fastq files before sequence alignment. Concatenated samples showed no irregularities in downstream analyses such as in elevated levels of autosomal heterozygosity, and were distributed across the species complex and across the DAPC clusters (Table S1).

Genotyping of the 711 bp region of the mitochondrial cytochrome oxidase subunit 1 (CO1) gene was conducted as per (Bierman et al. 2022) using the primer pair LCO1490 and HCO2198 (Leray et al. 2013). Briefly, DNA was extracted from beetle head and thorax tissue using Qiagen DNEasy Blood & Tissue kit protocols for insects (Purification of total DNA from insects (DY14 Aug-06; p 2–3; Qiagen (cat. no. 69504 or 69506)). PCR amplification was conducted on diluted DNA extracts (1:20 for quantities measured between 0.7 and 135 ng/µL), and 5 µL of the diluted DNA was used for PCR. AccuStart II PCR Master Mix (QuantBio) was used for the PCR reaction and conditions followed: 3 min 94 °C; 40 cycles of 30 s 94 °C, 30 s 50 °C, 1 min 72 °C. Agarose gel electrophoresis (1.5% agarose and SmartGlow Loading dye (Accuris Reagents)) was used to verify the amplification and presence of single fragments. These were used for Sanger sequencing in both directions at the Central Analytical Facility at Stellenbosch University. Samples were assigned to haplotypes described by (Stouthamer et al. 2017) by aligning sequences using BLAST (Camacho et al. 2009) to accession numbers from (Stouthamer et al. 2017).

### Sequence alignment

Raw genomic sequence reads were processed using Cutadapt v3.4 (Martin 2011) to remove adapters and Trimmomatic v0.39 (Bolger et al. 2014) to trim sequences to 100 bp. Sequences were aligned end-to-end to the PSHB reference assembly (Bickerstaff et al. 2024) using Bowtie2 v2.3.4.3 with --very-sensitive settings (Langmead and Salzberg 2012). Sam files were converted to bam and sorted in Samtools v1.7 (Danecek et al. 2021), read groups were added and validated with Picard v2.27.4 (Anon 2019), then Samtools was used to index bam files and filter unmapped reads and those not of primary alignment.

*Wolbachia* infection was assessed through alignment to the *w*Ei genome assembled in Bickerstaff et al. (2024) using identical settings to above. Samples were called as infected from their higher alignment rates relative to other samples, generally 10^2^–10^3^ times higher.

Initial analyses of genetic structure identified a series of sites distributed across the genome where heterozygosity was high across multiple individuals from different sampling locations. As truly heterozygous sites are likely to be caused by recent outbreeding, we deemed it unlikely that the same patterns would emerge in different locations. We identified 738 SNPs where 3 or more individuals were heterozygotes and these were from two or more sampling locations (e.g. KwaZulu-Natal and Western Cape; Fig. 1c). Although some of these sites had higher depth of coverage, pointing to them being caused by paralogs (McKinney et al. 2017), others did not have this pattern. As we were unable to determine what these sites represented, we filtered all RADtags (n=401) that contained one or more of these SNPs.

### Analytical pipelines for genetic structure and missense mutations

We ran the following pipeline to obtain and filter genotypes for discriminant analysis of principal components (DAPC (Jombart et al. 2010)), PCA, pairwise F_ST_, and private alleles.

Filtered bam files were built into Stacks catalogs using Stacks v2.65 program “ref_map” (Rochette et al. 2019) and VCFs were generated using the program “populations”. For analyses across the species complex, we retained SNPs scored in ≥0.9 of samples and ≥0.25 of samples from each population, and with minor allele count ≥3 to best ensure population-specific alleles were retained (where n=2 for China, North Vietnam, Florida and San Diego). For analyses within PSHB, we retained SNPs scored in ≥0.75 of samples and ≥0.25 of samples from each population and with a minor allele count ≥5. For DAPC and PCA, we also imputed genotypes using Beagle v4.0 (Browning and Browning 2016) in windows of 30 SNPs with a 10 SNP overlap.

DAPC was run in R package adegenet (Jombart 2008) using find.clusters to infer an optimum partitioning of the dataset into K clusters, then plotted using K-1 principal components to avoid overfitting (Thia 2023). Including the 401 RADtags with highly heterozygous SNPs did not affect downstream DAPC cluster analysis. Increasing K above 7 did not affect the inference of the hybrid clusters, but split the hybrid clusters into fewer clusters. PCA was run using R package LEA v3.12.2 (Gain and François 2021). Private allele analysis of population trios was run in “populations”, which was also used to estimate pairwise F_ST_.

The distribution of missense mutations was assessed with the Ensembl Variant Effect Predictor (McLaren et al. 2016), setting ‘minimal’ and ‘pick’ options. VEP inferred which variants represented synonymous substitutions and which were nonsynonymous substitutions. Stop codon mutations were ignored as these could be in pseudogenes or duplicate regions which are common in the PSHB genome (Bickerstaff et al. 2024).

### Analytical pipelines for D_XY_ and heterozygosity

We ran the following pipeline to obtain and filter genotypes for D_XY_. This pipeline was optimised to produce VCFs that retained both monomorphic and polymorphic sites (Korunes and Samuk 2021; Schmidt et al. 2024).

Filtered bam files were first processed using GATK v4.2.6.1 (Auwera and O’Connor 2020). GATK’s HaplotypeCaller was first used to generate gVCFs, set to EMIT_ALL_CONFIDENT_SITES. GenomicsDBImport imported gVCFs into a database, and they were genotyped together using GenotypeGVCFs set to --include-non-variant-sites. The genotyped VCF was indexed, indels were removed with SelectVariants, hard filtering was performed with VariantFiltration (D < 2.0, QUAL < 30.0, SOR > 3.0, FS > 60.0, MQ < 40.0), then SelectVariants was used to --set-filtered-gt-to-nocall.

For estimating D_XY_, we used pixy v1.2.5 (Korunes and Samuk 2021). We first used VCFtools to retain sites with sequencing depth ≥ 15. We ran pixy with a window size of 100,000, omitting the hybrid lineages in South Africa. Divergence time estimates were obtained from the formula for a strictly allopatric model (Nei 1987), where the expected value of D_XY_ = 2µt + π_Anc_, with µ the mutation rate, t the time since divergence, and π_Anc_ the level of diversity in the ancestral population. For µ we used a single-nucleotide mutation rate estimate for Colorado potato beetle of 5.8 × 10^-9^ (Xu et al. 2026). For π_Anc_, we reran pixy to estimate π across the native range lineages of Central Vietnam A, and used this estimate for all split times to allow for consistent comparisons. For all D_XY_ and π estimates, we omitted windows with fewer than 100 sites retained after filtering.

Bootstrap estimates of divergence times were obtained by subsampling with replacement 100 times from the set of windows retained after filtering.

For estimating observed heterozygosity, we first attempted to obtain estimates at both monomorphic and polymorphic sites using the above pipeline. However, we found that there were differences in the distribution of singletons between DArTseq runs, which strongly biased our estimates. Accordingly, we investigated heterozygosity only at SNPs with MAC≥3. While this would not provide estimates of autosomal heterozygosity (Schmidt, Jasper, et al. 2021), it was sufficient for identifying recent hybrids.

## Acknowledgments

We thank James Bickerstaff for providing reference fastas for *w*Ei and PSHB mtDNA and for useful discussions around PSHB biology. We thank Nathan Lo, Ros Gloag, and Maxim Adams for feedback on an earlier manuscript. For sampling assistance we thank Brett Hurley, Steffan Hansen, HortGro, Minette Karsten, Nanike Esterhuizen, Paul Barker, Shawn Fell, Trudy Paap, Wilhelm de Beer, You Li, Jiri Hulcr and Aaron Weber. For logistical support we thank Adriaan Engelbrecht and his group from UWC. We thank Marianne Coquilleau for illustrations of borers and brood galleries.

For funding we thank the Australian Research Council DE230100257 (TLS), Stellenbosch University Centre for Invasion Biology (JST, AB), the National Research Foundation (NRF) Foundational Biodiversity Information Programme (FBIP) (JST, AB), Stellenbosch Theoretical Institute for Advances Studies (STIAS) (JST), and NRF postdoctoral fellowship (EO).

## Author contributions

Conceptualization: TLS, AB, EJH, JST & AAH

Fieldwork: AB, EJH

Analysis: TLS

Visualization: TLS

Supervision: JST, AAH

Writing—original draft: TLS

Writing—review & editing: TLS, AB, EJH, JST & AAH

## Competing interests

The authors declare they have no competing interests.

## Data, code, and materials availability

Raw sequence data for all 247 shot-hole borer individuals are accessible through the NCBI SRA, BioProject ID PRJNA1508256. All metadata are listed in Table S1.

## Notes

### Competing Interest Statement

The authors have declared no competing interest.

### Summary of Updates

Has been peer reviewed, and substantial edits have been made to support the original findings.

## REFERENCES

Acer S, Hızal E, Ercan F. 2025. The First Report of Euwallacea fornicatus (Eichhoff, 1868) (Coleoptera: Scolytinae) in Türkiye with New Reproductive Host Plant. FRSTST 75:1–7.

Anon. 2019. Picard Toolkit. Available from: https://broadinstitute.github.io/picard/

Auwera GAV der, O’Connor BD. 2020. Genomics in the Cloud: Using Docker, GATK, and WDL in Terra. 1st ed. O’Reilly Media, Inc.

Bertorelle G, Raffini F, Bosse M, Bortoluzzi C, Iannucci A, Trucchi E, Morales HE, van Oosterhout C. 2022. Genetic load: genomic estimates and applications in non-model animals. Nat Rev Genet 23:492–503.

Bickerstaff JRM, Walsh T, Court L, Pandey G, Ireland K, Cousins D, Caron V, Wallenius T, Slipinski A, Rane R, et al. 2024. Chromosome structural rearrangements in invasive haplodiploid ambrosia beetles revealed by the genomes of *Euwallacea fornicatus* (Eichhoff) and *Euwallacea similis* (Ferrari) (Coleoptera, Curculionidae, Scolytinae). Genome Biol Evol 16:evae226.

Bierman A, Roets F, Terblanche JS. 2022. Population structure of the invasive ambrosia beetle, *Euwallacea fornicatus*, indicates multiple introductions into South Africa. Biol Invasions 24:2301–2312.

Bolger AM, Lohse M, Usadel B. 2014. Trimmomatic: a flexible trimmer for Illumina sequence data. Bioinformatics 30:2114–2120.

Browning BL, Browning SR. 2016. Genotype imputation with millions of reference samples. Am J Hum Genet 98:116–126.

Camacho C, Coulouris G, Avagyan V, Ma N, Papadopoulos J, Bealer K, Madden TL. 2009. BLAST+: architecture and applications. BMC Bioinformatics 10:421.

Carrillo D, Cruz LF, Kendra PE, Narvaez TI, Montgomery WS, Monterroso A, De Grave C, Cooperband MF. 2016. Distribution, pest status and fungal associates of *Euwallacea* nr. *fornicatus* in Florida avocado groves. Insects 7:55.

Ceriani-Nakamurakare E, Gomez DF, Trebino A, Listre A, Ingaramo L, Pilón AA, Bollazzi M. 2025. Increasing breeding host range and fast spread across Uruguay reveals the invasion potential of *Euwallacea fornicatus* (Coleoptera, Scolytinae) in South America. NeoBiota 98:247–260.

Ceriani-Nakamurakare E, Johnson AJ, Gomez DF. 2023. Uncharted Territories: First report of *Euwallacea fornicatus* (Eichhoff) in South America with new reproductive hosts records. Zootaxa 5325:289–297.

Charlesworth B, Charlesworth D. 1997. Rapid fixation of deleterious alleles can be caused by Muller’s ratchet. Genet Res 70:63–73.

Cooperband MF, Stouthamer R, Carrillo D, Eskalen A, Thibault T, Cossé AA, Castrillo LA, Vandenberg JD, Rugman-Jones PF. 2016. Biology of two members of the *Euwallacea fornicatus* species complex (Coleoptera: Curculionidae: Scolytinae), recently invasive in the U.S.A., reared on an ambrosia beetle artificial diet. Agric For Entomol 18:223–237.

Covre LS, Atkinson TH, Johnson AJ, Flechtmann CAH. 2024. Introduction and establishment of *Euwallacea fornicatus* (Coleoptera: Curculionidae: Scolytinae) in Brazil. J Econ Entomol 117:1192–1197.

Cunningham CB, Ji L, Wiberg RAW, Shelton J, McKinney EC, Parker DJ, Meagher RB, Benowitz KM, Roy-Zokan EM, Ritchie MG, et al. 2015. The genome and methylome of a beetle with complex social behavior, *Nicrophorus vespilloides* (Coleoptera: Silphidae). Genome Biol Evol 7:3383–3396.

Danecek P, Bonfield JK, Liddle J, Marshall J, Ohan V, Pollard MO, Whitwham A, Keane T, McCarthy SA, Davies RM, et al. 2021. Twelve years of SAMtools and BCFtools. GigaScience 10:giab008.

Eatough Jones M, Paine TD. 2015. Effect of Chipping and Solarization on Emergence and Boring Activity of a Recently Introduced Ambrosia Beetle (*Euwallacea* sp., Coleoptera: Curculionidae: Scolytinae) in Southern California. J Econ Entomol 108:1852–1859.

Eskalen A, Gonzalez A, Wang DH, Twizeyimana M, Mayorquin JS, Lynch SC. 2012. First report of a Fusarium sp. and its vector tea shot hole borer (*Euwallacea fornicatus*) causing Fusarium dieback on avocado in California. Plant Dis 96:1070–1070.

Facon B, Hufbauer RA, Tayeh A, Loiseau A, Lombaert E, Vitalis R, Guillemaud T, Lundgren JG, Estoup A. 2011. Inbreeding depression is purged in the invasive insect *Harmonia axyridis*. Curr Biol 21:424–427.

Figuet E, Nabholz B, Bonneau M, Mas Carrio E, Nadachowska-Brzyska K, Ellegren H, Galtier N. 2016. Life history traits, protein evolution, and the nearly neutral theory in amniotes. Mol Biol Evol 33:1517–1527.

Gain C, François O. 2021. LEA 3: Factor models in population genetics and ecological genomics with R. Mol Ecol Resour 21:2738–2748.

Glémin S. 2003. How are deleterious mutations purged? Drift versus nonrandom mating. Evol 57:2678–2687.

Goldarazena A, Alcazar-Alba MD, Hulcr J, Johnson AJ. 2025. First record of *Euwallacea fornicatus* Eichhoff (Coleoptera: Curculionidae: Scolytinae) in Spain. EPPO Bulletin 55:146–150.

Gomez DF, Johnson AJ. 2019. *Euwallacea fornicatus* (polyphagous shot-hole borer). CABI Available from: https://www.cabidigitallibrary.org/doi/10.1079/cabicompendium.18360453

Hoffmann AA, Turelli M, Harshman LG. 1990. Factors affecting the distribution of cytoplasmic incompatibility in *Drosophila simulans*. Genetics 126:933–948.

Hulcr J, Black A, Prior K, Chen C-Y, Li H-F. 2017. Studies of Ambrosia Beetles (Coleoptera: Curculionidae) in Their Native Ranges Help Predict Invasion Impact. flen 100:257–261.

Jombart T. 2008. adegenet: a R package for the multivariate analysis of genetic markers. Bioinformatics 24:1403–1405.

Jombart T, Devillard S, Balloux F. 2010. Discriminant analysis of principal components: a new method for the analysis of genetically structured populations. BMC Genetics 11:94–108.

Jordal BH, Normark BB, Farrell BD, Kirkendall LR. 2002. Extraordinary haplotype diversity in haplodiploid inbreeders: phylogenetics and evolution of the bark beetle genus *Coccotrypes*. Mol Phylogenet Evol 23:171–188.

Kalshoven LGE. 1958. Studies on the biology of Indonesian Scolytoidea. Entomol Ber 18:147–160.

Kilian A, Wenzl P, Huttner E, Carling J, Xia L, Blois H, Caig V, Heller-Uszynska K, Jaccoud D, Hopper C, et al. 2012. Diversity Arrays Technology: A Generic Genome Profiling Technology on Open Platforms. In: Pompanon F, Bonin A, editors. Data Production and Analysis in Population Genomics: Methods and Protocols. Totowa, NJ: Humana Press. p. 67–89.

Korunes KL, Samuk K. 2021. pixy: Unbiased estimation of nucleotide diversity and divergence in the presence of missing data. Mol Ecol Resour 21:1359–1368.

Lande R. 1994. Risk of Population Extinction from Fixation of New Deleterious Mutations. Evol 48:1460–1469.

Langmead B, Salzberg SL. 2012. Fast gapped-read alignment with Bowtie 2. Nat Methods 9:357–359.

Lantschner MV, Ceriani-Nakamurakare E, Johnson AJ, Cognato AI, Smith SM, Gomez DF. 2026. Invasion history reconstruction and potential distribution of the ambrosia beetles *Euwallacea fornicatus* and *E. perbrevis*, two global emerging pests. J Pest Sci 99:100.

Leray M, Yang JY, Meyer CP, Mills SC, Agudelo N, Ranwez V, Boehm JT, Machida RJ. 2013. A new versatile primer set targeting a short fragment of the mitochondrial COI region for metabarcoding metazoan diversity: application for characterizing coral reef fish gut contents. Front Zool 10:34.

Lockwood JL, Cassey P, Blackburn T. 2005. The role of propagule pressure in explaining species invasions. Trends Ecol Evol 20:223–228.

Lombaert E, Blin A, Porro B, Guillemaud T, Bernal JS, Chang G, Kirichenko N, Sappington TW, Toepfer S, Deleury E. 2025. Unraveling genetic load dynamics during biological invasion: insights from two invasive insect species. Peer Community J 5.

Lombaert E, Guillemaud T, Cornuet J-M, Malausa T, Facon B, Estoup A. 2010. Bridgehead Effect in the Worldwide Invasion of the Biocontrol Harlequin Ladybird. PLOS ONE 5:e9743.

Martin M. 2011. Cutadapt removes adapter sequences from high-throughput sequencing reads. EMBnet 17:10–12.

McKinney GJ, Waples RK, Seeb LW, Seeb JE. 2017. Paralogs are revealed by proportion of heterozygotes and deviations in read ratios in genotyping-by-sequencing data from natural populations. Mol Ecol Resour 17:656–669.

McLaren W, Gil L, Hunt SE, Riat HS, Ritchie GRS, Thormann A, Flicek P, Cunningham F. 2016. The Ensembl Variant Effect Predictor. Genome Biol 17:122.

Mendel Z, Lynch SC, Eskalen A, Protasov A, Maymon M, Freeman S. 2021. What Determines Host Range and Reproductive Performance of an Invasive Ambrosia Beetle *Euwallacea fornicatus*; Lessons From Israel and California. Front. For. Glob. Change 4.

Mendel Z, Protasov A, Maoz Y, Maymon M, Miller G, Elazar M, Freeman S. 2017. The role of *Euwallacea* nr. *fornicatus* (Coleoptera: Scolytinae) in the wilt syndrome of avocado trees in Israel. Phytoparasitica 45:341–359.

Mendel Z, Protasov A, Sharon M, Zveibil A, Yehuda SB, O’Donnell K, Rabaglia R, Wysoki M, Freeman S. 2012. An Asian ambrosia beetle *Euwallacea fornicatus* and its novel symbiotic fungus *Fusarium* sp. pose a serious threat to the Israeli avocado industry. Phytoparasitica 40:235–238.

Moir M, Kehoe M, Wright D, Bertazzoni S, Scott P, Wang C, Kinnaird E, Coutts B. 2025. The first Australian co-invasion of *Euwallacea fornicatus*, *Fusarium* sp. [AF18] and *Graphium euwallaceae*. BioInvasions Rec 14:575–585.

Nei M. 1987. Molecular Evolutionary Genetics. Columbia University Press

Nowell Nicolle L, Fournier-Level A, Robin C, Phillips BL. 2026. Genetic Allee Effects for Controlling Invasive Populations. Mol Ecol 35:e70228.

Paap T, de Beer ZW, Migliorini D, Nel WJ, Wingfield MJ. 2018. The polyphagous shot hole borer (PSHB) and its fungal symbiont *Fusarium euwallaceae*: a new invasion in South Africa. Australas Plant Pathol 47:231–237.

Peischl S, Dupanloup I, Kirkpatrick M, Excoffier L. 2013. On the accumulation of deleterious mutations during range expansions. Mol Ecol 22:5972–5982.

Peischl S, Kirkpatrick M, Excoffier L. 2015. Expansion Load and the Evolutionary Dynamics of a Species Range. Am Nat 185:E81–E93.

Phillips B. 2025. The ecology and evolution of invasive populations. Oxford University Press

Rochette NC, Rivera-Colón AG, Catchen JM. 2019. Stacks 2: Analytical methods for paired-end sequencing improve RADseq-based population genomics. Mol Ecol 28:4737–4754.

Rugman-Jones PF, Au M, Ebrahimi V, Eskalen A, Gillett CPDT, Honsberger D, Husein D, Wright MG, Yousuf F, Stouthamer R. 2020. One becomes two: second species of the *Euwallacea fornicatus* (Coleoptera: Curculionidae: Scolytinae) species complex is established on two Hawaiian Islands. PeerJ 8:e9987.

Rugman-Jones PF, Dodge CE, Stouthamer R. 2024. Pervasive heteroplasmy in an invasive ambrosia beetle (Scolytinae) in southern California. Heredity 133:388–399.

Ruzzier E, Ortis G, Vallotto D, Faccoli M, Martinez-Sañudo I, Marchioro M. 2023. The first full host plant dataset of Curculionidae Scolytinae of the world: tribe Xyleborini LeConte, 1876. Sci Data 10:166.

Schmidt TL, Endersby-Harshman N, Mills T, Rane R, Pandey G, Hardy C, Court L, Webb C, Trewin B, Neilan B, et al. 2025. Populations of the Australian Saltmarsh Mosquito *Aedes vigilax* Vary Between Panmixia and Temporally Stable Local Genetic Structure. Evol Appl 18:e70119.

Schmidt TL, Endersby-Harshman NM, Hoffmann AA. 2021. Improving mosquito control strategies with population genomics. Trends Parasitol 37:907–921.

Schmidt TL, Endersby-Harshman NM, Kurucz N, Pettit W, Krause VL, Ehlers G, Muzari MO, Currie BJ, Hoffmann AA. 2023. Genomic databanks provide robust assessment of invasive mosquito movement pathways and cryptic establishment. Biol Invasions 25:3453–3469.

Schmidt TL, Jasper M-E, Weeks AR, Hoffmann AA. 2021. Unbiased population heterozygosity estimates from genome-wide sequence data. Meth Ecol Evol 12:1888–1898.

Schmidt TL, van Rooyen AR, Chung J, Endersby-Harshman NM, Griffin PC, Sly A, Hoffmann AA, Weeks AR, Endersby-Harshman NM, Griffin PC, et al. 2019. Tracking genetic invasions: Genome-wide single nucleotide polymorphisms reveal the source of pyrethroid-resistant Aedes aegypti (yellow fever mosquito) incursions at international ports. Evol Appl 12:1136–1146.

Schmidt TL, Thia JA, Hoffmann AA. 2024. How can genomics help or hinder wildlife conservation? Annu Rev Anim Biosci 12.

Schwartz JH. 2021. Evolution, systematics, and the unnatural history of mitochondrial DNA. Mitochondrial DNA Part A 32:126–151.

Sendrowski J, Bataillon T. 2024. fastDFE: Fast and Flexible Inference of the Distribution of Fitness Effects. Mol Biol Evol 41:msae070.

Smith AL, Hodkinson TR, Villellas J, Catford JA, Csergő AM, Blomberg SP, Crone EE, Ehrlén J, Garcia MB, Laine A-L, et al. 2020. Global gene flow releases invasive plants from environmental constraints on genetic diversity. Proc Natl Acad Sci U S A 117:4218–4227.

Stouthamer R, Rugman-Jones P, Thu PQ, Eskalen A, Thibault T, Hulcr J, Wang L-J, Jordal BH, Chen C-Y, Cooperband M, et al. 2017. Tracing the origin of a cryptic invader: phylogeography of the *Euwallacea fornicatus* (Coleoptera: Curculionidae: Scolytinae) species complex. Agric For Entomol 19:366–375.

Subramanian S. 2013. Significance of Population Size on the Fixation of Nonsynonymous Mutations in Genes Under Varying Levels of Selection Pressure. Genetics 193:995–1002.

Sun J-T, Wang M-M, Zhang Y-K, Chapuis M-P, Jiang X-Y, Hu G, Yang X-M, Ge C, Xue X-F, Hong X-Y. 2015. Evidence for high dispersal ability and mito-nuclear discordance in the small brown planthopper, *Laodelphax striatellus*. Sci Rep 5:8045.

Thia JA. 2023. Guidelines for standardizing the application of discriminant analysis of principal components to genotype data. Mol Ecol Resour 23:523–538.

Tierney SM, Cooper SJB, Saint KM, Bertozzi T, Hyde J, Humphreys WF, Austin AD. 2015. Opsin transcripts of predatory diving beetles: a comparison of surface and subterranean photic niches. R Soc Open Sci. 2:140386.

Umeda C, Paine T. 2019. Temperature can limit the invasion range of the ambrosia beetle *Euwallacea* nr. *fornicatus*. Agric For Entomol 21:1–7.

Warren RJ, Mokadam C. 2025. Asexuality and species invasion. Biodivers Conserv 34:29–43.

Weber CC, Nabholz B, Romiguier J, Ellegren H. 2014. Kr/Kc but not dN/dS correlates positively with body mass in birds, raising implications for inferring lineage-specific selection. Genome Biol 15:542.

de Wit MP, Crookes DJ, Blignaut JN, de Beer ZW, Paap T, Roets F, van der Merwe C, van Wilgen BW, Richardson DM. 2022. An Assessment of the Potential Economic Impacts of the Invasive Polyphagous Shot Hole Borer (Coleoptera: Curculionidae) in South Africa. J Econ Entomol 115:1076–1086.

Xu S, Al-Madhagy S, Duchen P, Edison A. 2026. Trio-sequencing Reveals High Germline Mutation Rates in the Colorado Potato Beetle (*Leptinotarsa decemlineata*). Genome Biol Evol 18:evag027.

