## Supplementary material for "Ambrosia beetle invasions are structured by inbreeding, intraspecific hybridisation, and bridgeheads": Figures S1-S4

**Fig. S1. PCAs of SHB.**

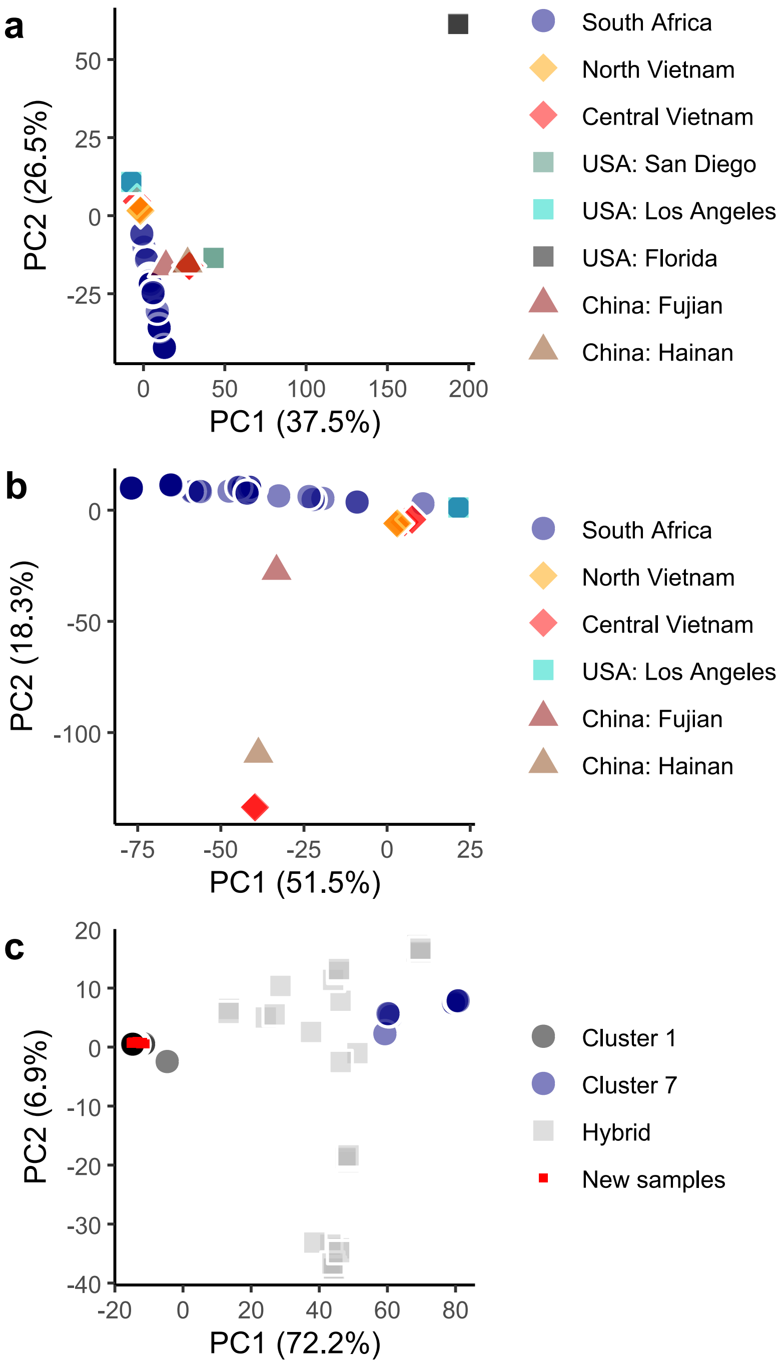


(a) PCA of all SHB populations. (b) PCA of all PSHB populations. (c) PCA of South African PSHB samples, including the 59 additional samples from 26 locations (Fig. 2c).

**Fig. S2. Heatmap showing PSHB genotypes across the first six chromosomes (I–VI).**

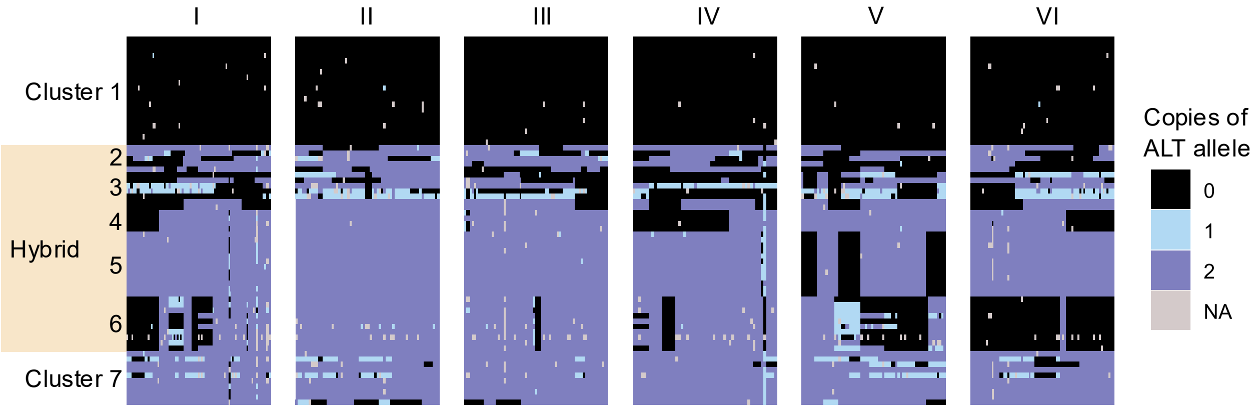


Individuals are sorted by cluster on y-axis. SNP genotypes are sorted by position on x-axis.

**Fig. S3. Proportion of alleles identical to the reference genome.**

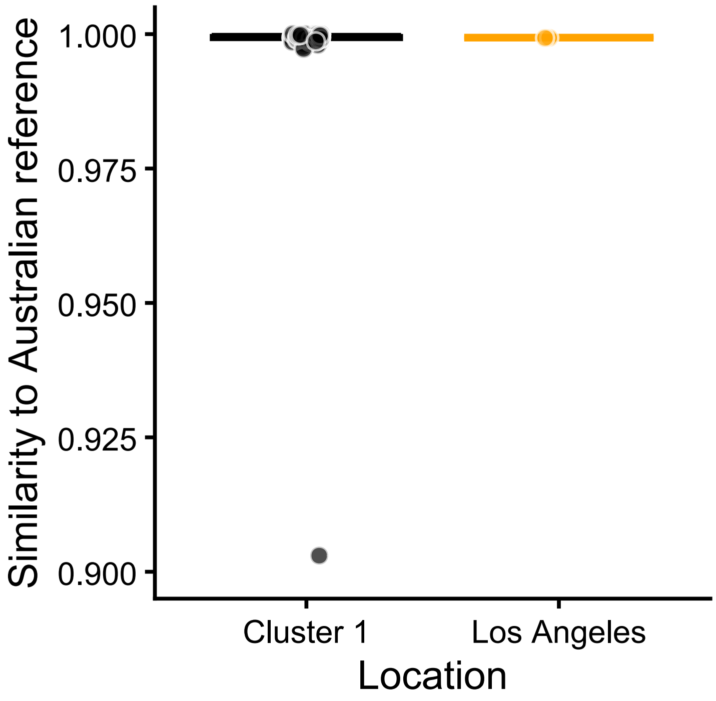


Alleles are at 3060 SNPs called within South African Cluster 1 and 7, and hybrid clusters. Thirteen Cluster 1 samples had 100% similarity, while one sample was admixed. Each Los Angeles sample had four non-reference alleles.

**Fig. S4. Distribution of Cluster 1 mtDNA haplotypes in South Africa**

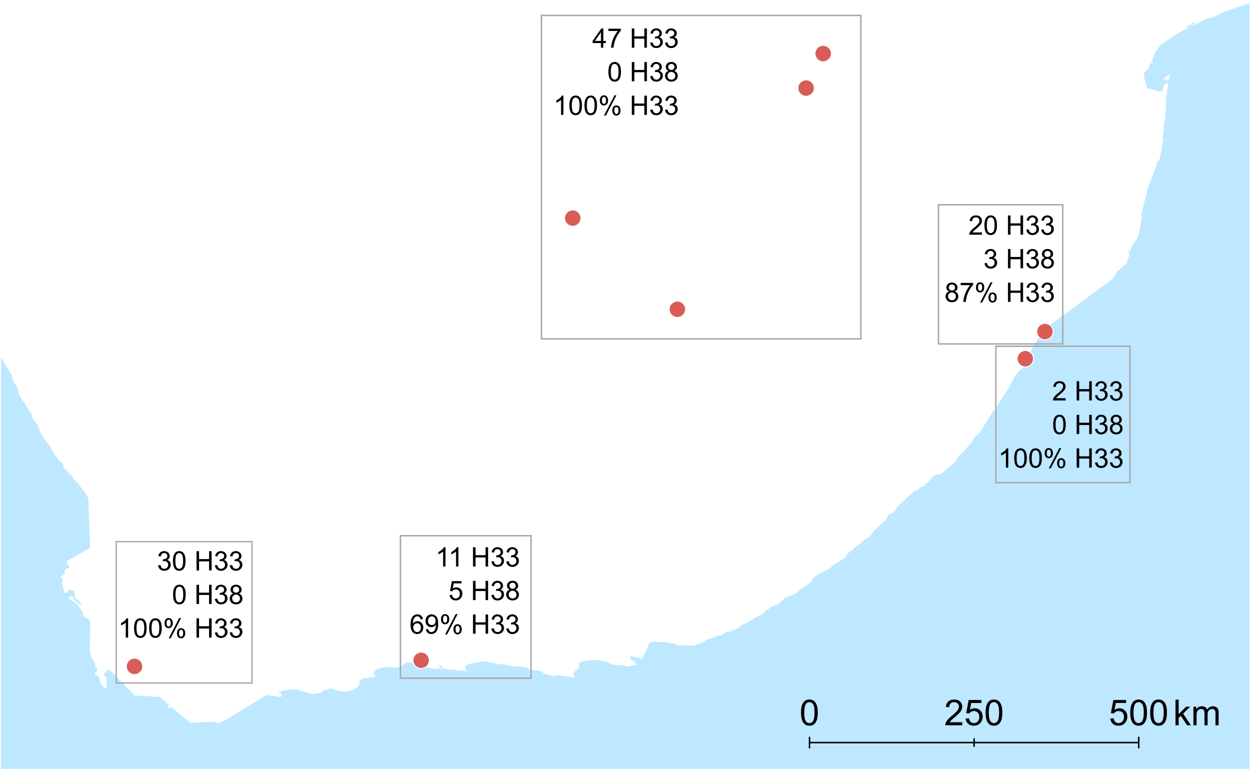


Symbols indicate broad sampling locations. Boxes summarise the local haplotype proportions. Haplotypes of Clusters 2–7 are in Table S1.
